# Plant Functional Type Composition, Rather Than Species Diversity, Shapes Soil Microbial Functional Diversity and Redundancy

**DOI:** 10.64898/2026.07.13.738212

**Authors:** Nichole Giani, Prabhsimran Singh, Vidya Suseela, Barbara Campbell

## Abstract

Cover crops (CCs) are widely used to improve soil health, but their species-specific effects on microbial functional diversity and redundancy remain poorly understood. We evaluated how monocultures of field pea (*Pisum sativum*), forage radish (*Raphanus sativus*), and cereal rye (*Secale cereale*), as well as a three-species mixture (3spp) and a five-species mixture (5spp), influence rhizosphere bacterial and fungal communities in a field experiment. Amplicon sequencing and predictive functional profiling were used to assess microbial diversity, composition, functional potential, and functional redundancy (FR). Microbial alpha diversity showed little change across treatments, but community composition and predicted functional profiles were strongly influenced by CC identity. The 3spp rhizobiome showed the highest bacterial FR, broad metabolic capacity, and greater network connectivity, suggesting increased functional stability of all treatments. In contrast, the 5spp rhizobiome showed reduced bacterial FR and more widespread functional depletion, likely due to imbalances in plant functional types. Monocultures showed species-specific patterns, with rye supporting functionally efficient communities and radish promoting more competitive interactions and reduced functional diversity. Fungal communities responded differently from bacterial communities. While fungal taxonomic shifts were limited, they showed stronger compositional differences among treatments and more stable functional profiles. Notably, fungal FR was highest in the 5spp rhizobiome. Overall, these findings highlight that balanced plant functional composition, rather than greater species richness alone, is important for shaping rhizosphere microbial function and redundancy. Balanced mixtures promoted higher bacterial redundancy, while more diverse but functionally imbalanced mixtures did not consistently enhance microbial function.

**Importance:** The benefits of using CCs in agricultural settings are mostly influenced by microbial communities, including nutrient cycling, organic matter turnover, and resilience to disturbance. However, CC mixtures are often promoted on the assumption that more species provide greater soil benefits. This work challenges that assumption by showing that the functional identity and balance of CC species may matter more than the number of species alone. This distinction is important for farmers, land managers, and researchers because CC seed mixes can be costly, and poorly balanced mixtures may fail to deliver intended microbial benefits. Understanding which plant combinations support functionally stable microbial communities can improve CC recommendations and help design agricultural systems that maintain soil processes under changing environmental conditions.

## Introduction

Soil microorganisms play a crucial role in maintaining soil health and crop productivity, as they facilitate nutrient cycling, break down organic matter, promote plant growth, and help the ecosystem recover from environmental stress (1). Microbial communities carry out a wide range of biochemical functions that drive carbon, nitrogen, phosphorus, and sulfur cycling, among others (2). Not only is the presence of a diverse microbial community important, but the degree of functional redundancy (FR), the number of organisms that can perform the same function, is also key to maintaining ecosystem stability (3, 4). FR has been proposed as a mechanism to buffer against microbial community collapse, because if one taxon is lost due to disturbance, another may still perform the same role, thereby maintaining overall ecosystem stability (5, 6).

In agriculture, the use of cover cropping to enhance microbial diversity and FR is gaining interest, particularly for long-term soil fertility and resilience to stress. Cover crops (CC) are grown between the cycles of cash crops. They are known to improve soil quality by preventing erosion, enhancing soil structure, increasing organic matter, suppressing weeds and pests, and improving nutrient availability (7, 8). CCs can significantly influence the composition and function of soil microbial communities through root exudation, organic matter inputs, and rhizosphere interactions (9, 10). However, relatively little is known about how individual CC species differ in their ability to enrich for specific microbial taxa and promote functional diversity and redundancy. Previous studies have demonstrated that CCs can significantly influence soil microbial communities, although the magnitude and direction of these effects often depend on species identity and diversity. For example, a global meta-analysis found that cover cropping increased microbial biomass and altered microbial community composition compared to fallow soils, with particularly strong responses observed in fungal communities (11).

Similarly, studies examining the decomposition of CC residues showed that mixtures with multiple species can increase microbial functional diversity relative to monocultures due to the broader range of substrates available to soil microbes (12). While previous research has shown that CCs generally increase microbial biomass, enzyme activity, and sometimes diversity (13, 14), fewer studies have examined how individual CC species affect FR. Most studies on soil microorganisms under different CCs focus on taxonomic diversity or community composition (11, 15, 16), with less emphasis on the stability and resilience conferred by overlapping microbial functions. For example, microbial alpha diversity had only subtle impacts on enzyme activity in cover-cropped soils, with total community size more closely associated with functions than with diversity, suggesting high FR (17). While theoretical work indicates that FR acts as a buffer against biodiversity loss in microbial communities (18), few studies have explicitly examined how individual CC species influence this redundancy and its effects on ecosystem stability (19, 20). This knowledge gap limits our ability to predict which CC species are most effective at promoting resilient microbial communities that can withstand environmental changes or disturbances.

Cover crop species differ in their physiology, nutrient use strategies, and root exudate profiles, all of which can influence the microbial communities in the surrounding soil. Legumes such as Austrian winter pea (*Pisum sativum*) host nitrogen-fixing bacteria and generally support communities enriched in taxa, including *Rhizobium* and *Bradyrhizobium* (21, 22). Grasses (e.g., oats; *Avena sativa L*.) and brassicas (forage radish; *Raphanus sativus L*.) may support different groups of microbes, including those involved in phosphorus solubilization, sulfur oxidation, or other nutrient pathways (9, 23, 24). Some of these interactions may be species-specific, because particular plant species select for microbial communities through root exudates that can act as carbon sources or chemical signals (25, 26). These plant-microbe interactions are dynamic and can influence both the taxonomic composition and functional capabilities of the soil microbiome.

In this study, we focused on determining how different CCs affect microbial taxonomic and predicted functional diversity and redundancy in field soils. Specifically, we hypothesize that microbial taxonomic diversity, functional diversity, and FR will differ among CC species, with legumes (e.g., pea) supporting higher redundancy due to their enrichment of beneficial microbes involved in nitrogen cycling and plant growth. We further hypothesize that CC mixtures will support greater microbial taxonomic and functional diversity, as well as increased FR, compared to monocultures, due to the greater diversity of plant inputs and ecological niches provided by CC mixtures. To test these hypotheses, we used 16S rRNA gene and ITS amplicon sequencing to assess microbial community composition and diversity and employed functional prediction tools to estimate functional potential and redundancy from these data. This approach will provide insight into the extent to which amplicon-based methods can reveal species-specific effects of CCs on soil microbial communities.

Overall, this work addresses key gaps in our understanding of how specific plant species may influence microbial functional stability in field soils. By applying amplicon sequencing and predictive functional profiling, we can identify which CC species are most likely to support resilient, functionally redundant microbial communities, with direct applications for improving agricultural soil management in the face of changing environmental conditions.

## Materials and Methods

### Crop Information and DNA Extraction

We established a CC cash crop rotation experiment in Fall 2023 at Simpson’s Farm in Pendleton, SC (34°37′30.1′′ N, −82°44′13.9′′ W). Cover crops were planted on November 8, 2023, using a no-till drill-cone plot planter. The six CC treatments were planted in the field in a randomized block design with four blocks and four replicates per treatment. There was a total of 24 plots, each 3 m x 15 m in size, with a 2m buffer zone between the plots and blocks. The soil at the site is Cecil sandy loam (clayey, kaolinitic, thermic typic Kanhapludults). The CCs were grown without any external irrigation or fertilizer inputs. Cover crops treatments were field pea (*Pisum sativum*), forage radish (*Raphanus sativus*), cereal rye (*Secale cereale*), a three-species mixture (3 spp.: field pea, oats (*Avena sativa*), forage radish), a five-species mixture (5 spp.: field pea, red clover (*Trifolium pratense*), crimson clover (*Trifolium incarnatum*), cereal rye, canola (*Brassica napus*)), and no CC fallow (control), which was fallow for six months prior to the start of the experiment. Seeding rates were based on local USDA recommendations (27). Cover crop harvest and soil sampling took place on March 26, 2024. Average temperatures ranged from 40.7 ℉ to 54.5 ℉ (average 48 ℉), and monthly precipitation ranged from 0.86 to 11 inches (average 6 inches) from November 2023 to March 2024. DNA was extracted from frozen rhizosphere using a modified phenol-chloroform-based method (28) and cleaned using the Zymo DNA Clean & Concentrator-5 kit and quantified using Qubit™ dsDNA HS assays (Thermo Fisher Scientific).

### Soil Analyses

Standard soil nutrient analysis was performed by the Clemson University Agricultural Service Lab. Three extracellular enzymes, namely, N-acetylglucosaminidase (NAG, EC 3.2.1.52), a nitrogen-acquiring enzyme; acid phosphatase (AP, EC 3.1.3.2), a phosphorus-acquiring enzyme; and β-glucosidase (BG, EC 3.2.1.21), a C-acquiring enzyme, were analyzed using the microplate assay technique (29, 30). Briefly, 5.0 g of defrosted soil was blended with 250 mL of 50 mM sodium acetate buffer (pH 6.0). 200 μL of soil slurry was combined with 50 μL of 200 μM methylumbelliferone (MUB) substrate in black 96-well microplate wells. The 96-well plates were incubated for 2 hours at room temperature in the dark, and fluorescence was measured spectrophotometrically at 365 nm (excitation) and 450 nm (emission). Enzyme activities were expressed as nmol h⁻¹ g⁻¹ soil. Finely ground samples were analyzed for total organic carbon and total nitrogen using a LECO CN analyzer (model: CN828). The low-carbon LECO soil standard (1.006 ± 0.20% C) was used to prepare the calibration curve.

### 16S rRNA gene and ITS Sequencing and Analysis

The V3-V4 region of the 16S rRNA was amplified using the dual-indexing strategy of Kozich et al. (2013) with primers 341F and 806R-B (31, 32). For ITS region one, primers ITS1F and ITS2 (33) were used with the same barcodes, linkers and Illumina adapters as in (32). PCR amplifications were performed using Q5 High-Fidelity DNA Polymerase (New England Biolabs) under the following conditions: initial denaturation at 95 °C for 2 min; 34 cycles of 95 °C for 20 s, 55 °C for 15 s, and 72 °C for 5 min; and a final extension at 72 °C for 10 min. Amplicons were purified with MagMAX™ Pure Bind Beads following the manufacturer’s protocol (Applied Biosystems). PCR products were quantified with Qubit assays, and Nanodrop spectrophotometry was used to measure DNA concentrations. A subset of samples quantified by both methods was used to generate a calibration curve for estimating Qubit-equivalent concentrations from Nanodrop values. Purified amplicons were normalized and pooled to 5 ng/µL, and the final pool was assessed on an Agilent TapeStation to confirm fragment size and calculate molarity. Sequencing was performed on an Illumina NextSeq 2000 with 600-cycle XLEAP chemistry (P2 flow cell for 16S rRNA gene amplicons, P1 flow cell for ITS amplicons).

### Sequence Processing and Bioinformatics

Paired-end sequences were processed with the Clemson University Palmetto Cluster using QIIME2 v.2022.11 (34). Reads were demultiplexed, quality-filtered, denoised, and merged using DADA2 (35), and chimeric sequences were removed. Representative Amplicon Sequence Variants (ASV) were aligned with MAFFT (36), and phylogenetic trees were constructed with FastTree2 (37) and midpoint-rooted. For diversity analyses, samples were rarefied to a depth of 334,132 reads per sample for the 16S rRNA gene and 3,477 for ITS sequences. Taxonomic assignments were performed with the q2-feature-classifier plugin using a Naïve Bayes classifier trained on SILVA 138 (16S rRNA gene) (38) or UNITE v9 (ITS1 region) (39). Resulting ASV tables, taxonomies, and metadata were imported into R (v.4.4.2) phyloseq (40) and microeco (41).

Alpha diversity metrics (Shannon, Richness, and Dominance (Berger-Parker) and Faith’s PD) were calculated with phyloseq (40, 42, 43). Statistical comparisons were conducted using ANOVA for normally distributed data and Kruskal–Wallis for non-parametric data, with Tukey’s HSD or pairwise Wilcoxon rank-sum tests (FDR-corrected) for post-hoc analyses (44–47). Beta diversity was evaluated with Bray–Curtis, Jaccard, and weighted/unweighted UniFrac distances, followed by PCoA ordination (48, 49). Group differences were tested with PERMANOVA (adonis2 package) (50) with pairwise comparisons adjusted for multiple testing. Homogeneity of dispersions was checked using betadisper (51). To evaluate whether technical replicates differed more than biological plot-level variation within the same CC treatment, pairwise beta-diversity distances were calculated among all samples using Bray–Curtis and UniFrac distances. Pairwise distances were then classified as either technical replicate pairs from the same plot, defined as samples with the same CC and plot number but different replicate IDs, or same-cover-crop different-plot pairs, defined as samples from the same We first evaluated whether variation among biological replicates exceeded variation between technical replicates, as one technical replicate was collected from each cover crop rhizobiome sampling plot (Supplemental Table 1). For both bacterial 16S and fungal ITS communities, Bray–Curtis distances between technical replicate pairs were not significantly different from distances between samples collected from different plots within the same cover crop treatment. For 16S communities, technical replicates had a slightly lower median Bray–Curtis distance than same-cover-crop, different-plot pairs, with median distances of 0.537 and 0.592, respectively. A similar pattern was observed for ITS communities, for which median Bray–Curtis distances were 0.636 for technical replicate pairs and 0.686 for same-cover-crop, different-plot pairs. These small differences indicate that technical replicates were slightly more similar to one another than biological samples from different plots receiving the same cover crop treatment, but technical replication did not introduce greater variability. Therefore, technical replicates were retained in downstream bacterial and fungal community analyses.v treatment but different plot numbers. Median distances were compared between these two groups to determine whether technical replicates were more dissimilar than independent plot samples from the same cover crop treatment.

### Differential Abundance and Functional Predictions

To identify differentially abundant taxa, ANCOM-BC2 (Lin & Peddada, 2020) was applied at multiple taxonomic ranks (Phylum, Class, Order, Family, Genus, and ASV) considering global and pairwise comparisons with FDR correction. For both 16S rRNA gene and ITS datasets, taxa were filtered prior to genus- and ASV-level ANCOM-BC2 analyses by retaining taxa present in at least 20% (16S) or 10% (ITS) of samples and with ≥10 total reads.

Functional predictions were generated from 16S rRNA gene ASV data using the PICRUSt2 database (52). Functional predictions were generated from ITS1 region sequences using the FungalTraits database (53). Microeco was used to prepare input microtable objects and calculate KEGG-based functional profiles (41). The FR Index (FRI) was computed to estimate redundancy within predicted pathways for bacterial communities (54). FR for fungal communities was calculated using the “SYNCSA” package in R based on the FungalTrait database (55–58). Global and pairwise differences in functional profiles and redundancy indices among treatments were tested using ANOVA or Kruskal–Wallis tests with appropriate post-hoc corrections (44, 45). To assess the reliability of functional predictions, weighted Nearest Sequenced Taxon Index (NSTI) scores were calculated for all samples using PICRUSt2 (59). NSTI represents the average phylogenetic distance between each ASV and its nearest reference genome, weighted by abundance. Lower NSTI values indicate higher prediction accuracy (59).

### Data Availability

16S rRNA gene and ITS sequences were uploaded to NCBI under the BioProject number PRJNA1416991; BioSample numbers SAMN54986586-SAMN54986615 (16S rRNA gene) and SAMN54986586-SAMN54986615 (ITS); and SRA numbers SRR38416195-SRR38416254.

## Results

### Rhizosphere Bacterial and Fungal Community Diversity in Response to Cover Crop Species

We first evaluated whether variation among biological replicates exceeded variation between technical replicates, as one technical replicate was collected from each CC rhizobiome sampling plot (Supplemental Table 1). For both bacterial 16S and fungal ITS communities, Bray–Curtis distances between technical replicate pairs were not significantly different from distances between samples collected from different plots within the same CC. For 16S communities, technical replicates had a slightly lower median Bray–Curtis distance than same-cover-crop, different-plot pairs, with median distances of 0.537 and 0.592, respectively. A similar pattern was observed for ITS communities, for which median Bray–Curtis distances were 0.636 for technical replicate pairs and 0.686 for same-cover-crop, different-plot pairs. These small differences indicate that technical replicates were slightly more similar to one another than biological samples from different plots receiving the same CC treatment, but technical replication did not introduce greater variability. Therefore, technical replicates were retained in downstream bacterial and fungal community analyses.

We assessed bacterial and fungal alpha and beta diversity to evaluate the effect of different CC species on rhizosphere or control communities (rhizobiomes). There were no significant differences for Shannon diversity, richness, or Faith’s PD for bacterial rhizobiomes (Figs. 1A, B, D). While not significant, the pea rhizobiome had the lowest Shannon diversity, richness and Faith’s PD. However, there were significant differences in dominance between 3 spp. vs pea and rye vs pea, with the pea bacterial rhizobiome having the highest dominance overall (Fig. 1C). There were also no significant differences in Shannon diversity, richness, or dominance in the fungal rhizobiomes (Figs. 1E, 1F, 1G) but there was a significant difference in Faith’s PD when comparing 5spp vs radish, with the 5spp fungal rhizobiome showing the highest phylogenetic diversity overall (Fig. 1H) The radish rhizobiome had low values of overall fungal diversity, richness and Faith’s PD, but a high value for dominance.

**Figure 1:**
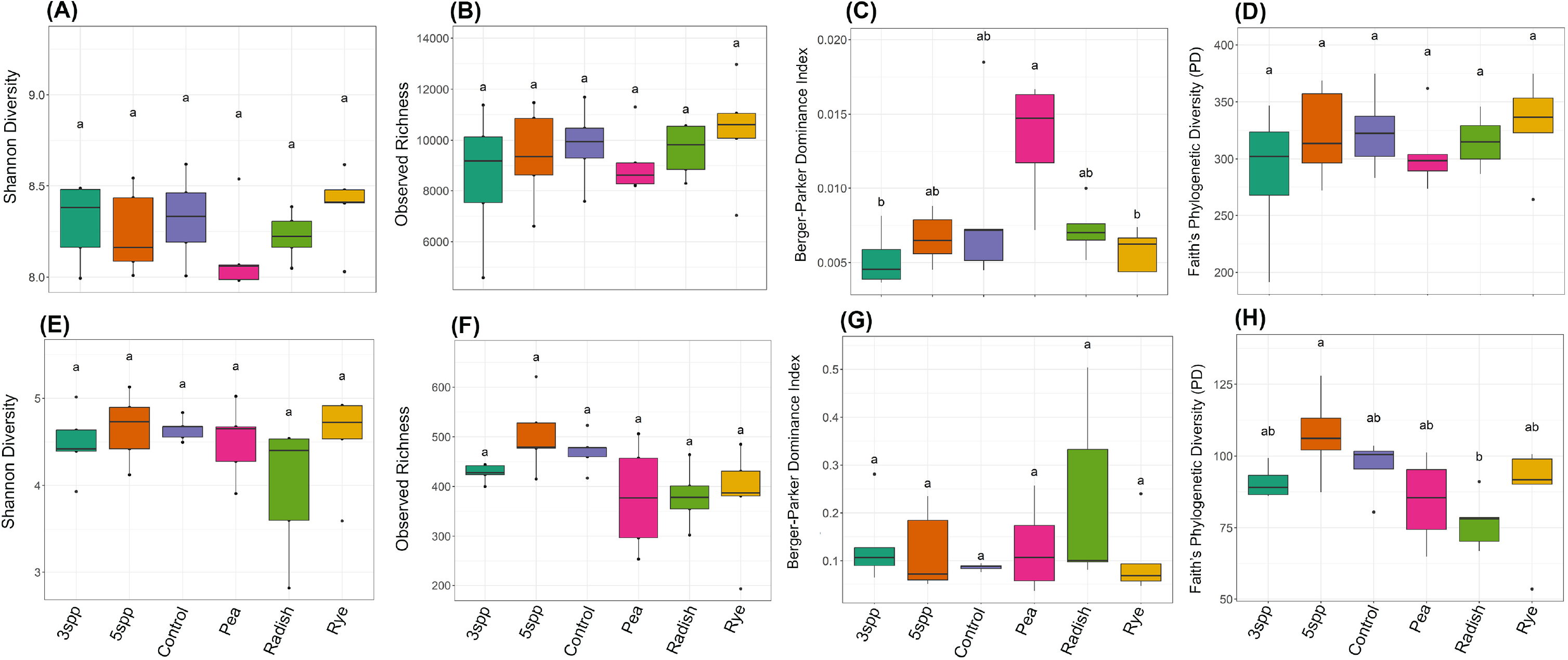
Alpha diversity of rhizosphere bacterial and fungal communities. Panels A–D show bacterial Shannon diversity, richness (Observed), dominance (Berger-Parker) and Faith’s PD, respectively. Panels E-H show fungal Shannon diversity, richness (Observed), dominance (Berger-Parker) and Faith’s PD, respectively. Different letters indicate statistically significant differences between groups (p ≤ 0.05).

Beta diversity analyses revealed separation in overall community composition for bacterial (p = 0.009; Fig. 2A) and fungal (p = 0.001; Fig. 2B) communities based on CC identity. Bacterial communities, however, did not exhibit any significant pairwise differences in beta diversity between CCs. In contrast, the rhizobiomes of several CCs exhibited significant pairwise differences among fungal communities, including 3spp. vs. radish, 5spp. vs. radish, 5spp. vs. rye, control vs. rye, and radish vs. rye.

**Figure 2.**
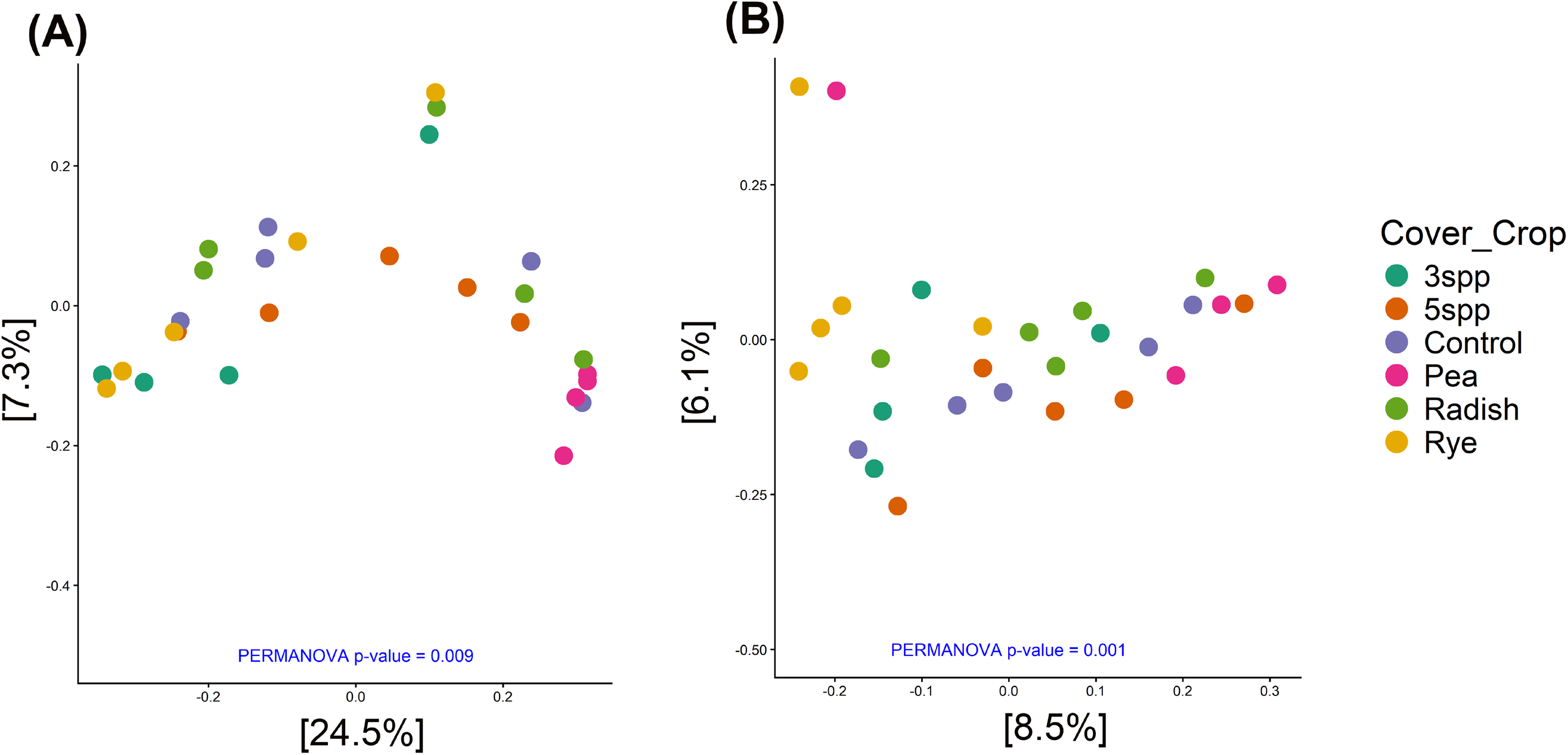
Principal Coordinates Analysis (PCoA) of bacterial (A) and fungal (B) communities. PCoA of bacterial and fungal communities based on Unifrac distance, with significance determined by PERMANOVA (p-values shown).

To determine if soil properties influenced the bacterial and fungal communities, the concentrations of various nutrients (phosphorus (P), potassium (K), calcium (Ca), magnesium (Mg), zinc (Zn), manganese (Mn), copper (Cu), boron (B), sodium (Na) and Nitrate-nitrogen (NO3-N)) in the rhizospheres of different CCs at the time of harvest. The only nutrients that were significantly different among CC soils were Mn and NO3-N (Supplemental Table 2). Mn was significantly different between 3spp and 5spp soils (p = 0.03). NO3_N was significantly different between 3spp vs control (p = 0.05), 5spp vs pea (p = 0.03), control vs pea (p = 0.003), control vs radish (p = 0.01) and pea vs rye (p = 0.05) soils. We also analyzed total carbon, total nitrogen, C:N ratio, and three different enzymes (Alkaline Phosphatase (AP), NG (N-5acetyl-β-d-glucosaminidase), BG (β -glucosidase)) and there were no significant differences among CCs for these variables (Supplemental Fig. 1).

Distance-based redundancy analysis (dbRDA) indicated that Mg and Ca concentrations and pH were significantly associated with bacterial community composition across CC treatments (overall model: Adjusted R² = 0.14; Adjusted R² based on CCs = 0.10; p = 0.016; Fig. 3A). Among these variables, Mg showed a significant association with community structure with higher Mg concentrations corresponding to samples from the 3spp rhizobiome. For fungal communities, pH and Ca were significantly associated with variation in community composition (overall model: Adjusted R² =0.077; Adjusted R² based on CCs = 0.061; p = 0.003; Fig. 3B). Soil pH was significant, with higher pH values corresponding to rye-associated rhizobiome samples. In contrast, total carbon, total nitrogen, and enzyme activities (acid phosphatase [AP], N-acetylglucosaminidase [NAG], and β-glucosidase [BG]) were not significantly associated with either bacterial or fungal community composition (Supplemental Fig. 1).

**Figure 3.**
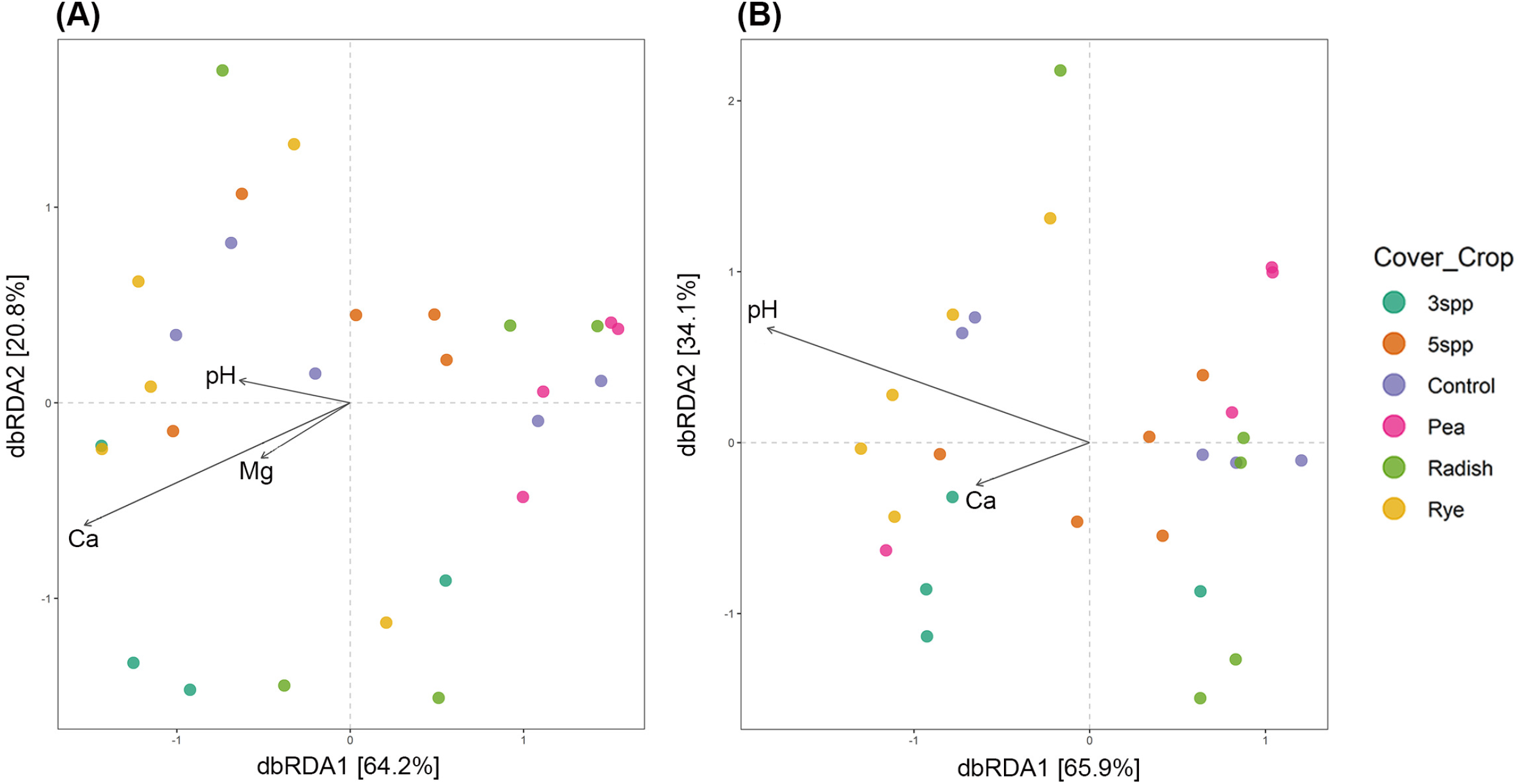
Distance-based Redundancy Analysis (dbRDA) of bacterial communities (A) and fungal communities (B) for soil pH and nutrients. Arrows indicate soil chemical variables associated with shifts in community composition. Only significant variables are shown.

Detailed taxonomic relative abundance patterns and all significant ANCOM-BC results are provided in Supplemental Fig. 2, Supplemental Data, and Supplemental Table 3. In brief, CC identity was associated with shifts in multiple bacterial taxa across taxonomic levels, including members of *Proteobacteria* (e.g., *Burkholderiaceae*), *Actinobacteriota* (e.g., *Frankiaceae*), and *Acidobacteriota*, which varied in relative abundance among treatments. For example, *Gammaproteobacteria* were more abundant in rye-associated rhizobiomes, while *Vicinamibacteria* were enriched in 3spp. Fungal communities showed significant beta diversity differences among treatments, but these compositional shifts were accompanied by relatively few differentially abundant taxa and no major changes in dominant higher-level lineages. Across treatments, fungal communities remained dominated by *Ascomycota*, particularly

### Sordariomycetes and Nectriaceae

Co-occurrence network complexity of bacterial communities varied among CCs (Supplemental Fig. 3 and Supplemental Table 4). Network complexity in the rhizobiomes, based on total number of edges, ranked as follows: 3spp > pea > 5spp > rye > control > radish. The 3spp and rye networks showed predominantly positive correlations, while 5spp, control, pea, and radish networks exhibited more negative correlations than the former. All rhizobiome networks were dominated by ASVs within four phyla (*Actinobacteriota*, *Planctomycetota*, *Proteobacteria*, *Chloroflexi*). *Actinobacteriota*-to-*Actinobacteriota* ASV connections were among the most abundant across all treatments, primarily positive in 3spp, but negative in all other CC rhizobiomes. Inter-phyla connections involving Chloroflexi ASVs showed variable patterns: predominantly negative with *Actinobacteriota* in most treatments, but positive with *Verrucomicrobiota* in 5spp. Fungal networks showed minimal complexity, with only three significant networks detected for 5spp, pea and rye all consisting of mostly positive interactions in pea and mostly negative interactions in 5spp and rye (Supplemental Fig. 3). The phylum with the most ASV interactions for all CCs was *Ascomycota*-to-*Ascomycota*.

### Differences in Bacterial and Fungal Predicted Functional Potential between samples

Weighted NSTI scores from PICRUSt2 ranged from 0.072 to 0.086 (mean ± SD: 0.078 ± 0.004) across all samples, indicating a moderate quality of functional predictions with adequate availability of closely related reference genomes (Supplemental Table 5). All samples fell below the conservative threshold of 0.15, suggesting acceptable reliability for downstream functional analyses. NSTI values did not differ substantially among treatments. PICRUSt2-based predictions of KEGG Category 2 functions revealed multiple significant differences among CC rhizobiomes (Fig. 4). At KEGG Category 2 (Fig. 4A), bacterial functional categories showed divergent responses to CC treatments. The 3spp. rhizobiome exhibited elevated scaled abundance in amino acid metabolism, carbohydrate metabolism, energy metabolism, and lipid metabolism, among others, compared to other CC rhizobiomes. While not as much as the 3spp rhizobiome, the radish rhizobiome showed enrichment across most metabolic categories, particularly in cell motility, glycan biosynthesis and metabolism, and transcription, while showing slight depletion in amino acid metabolism, environmental adaptation and xenobiotics biodegradation and metabolism. The rye rhizobiome showed increased abundance in environmental adaptation, xenobiotics biodegradation and metabolism, membrane transport and metabolism of other amino acids, but depletions in transcription and translation categories. The 5spp. showed the most significant depletion across most functional categories, with negative-scaled abundance values lower than those in the control and in most monoculture rhizobiomes. Except for a few categories, the pea rhizobiome mostly exhibited depletions compared to other CC, particularly enriched in cell motility, transcription and translation.

**Figure 4.**
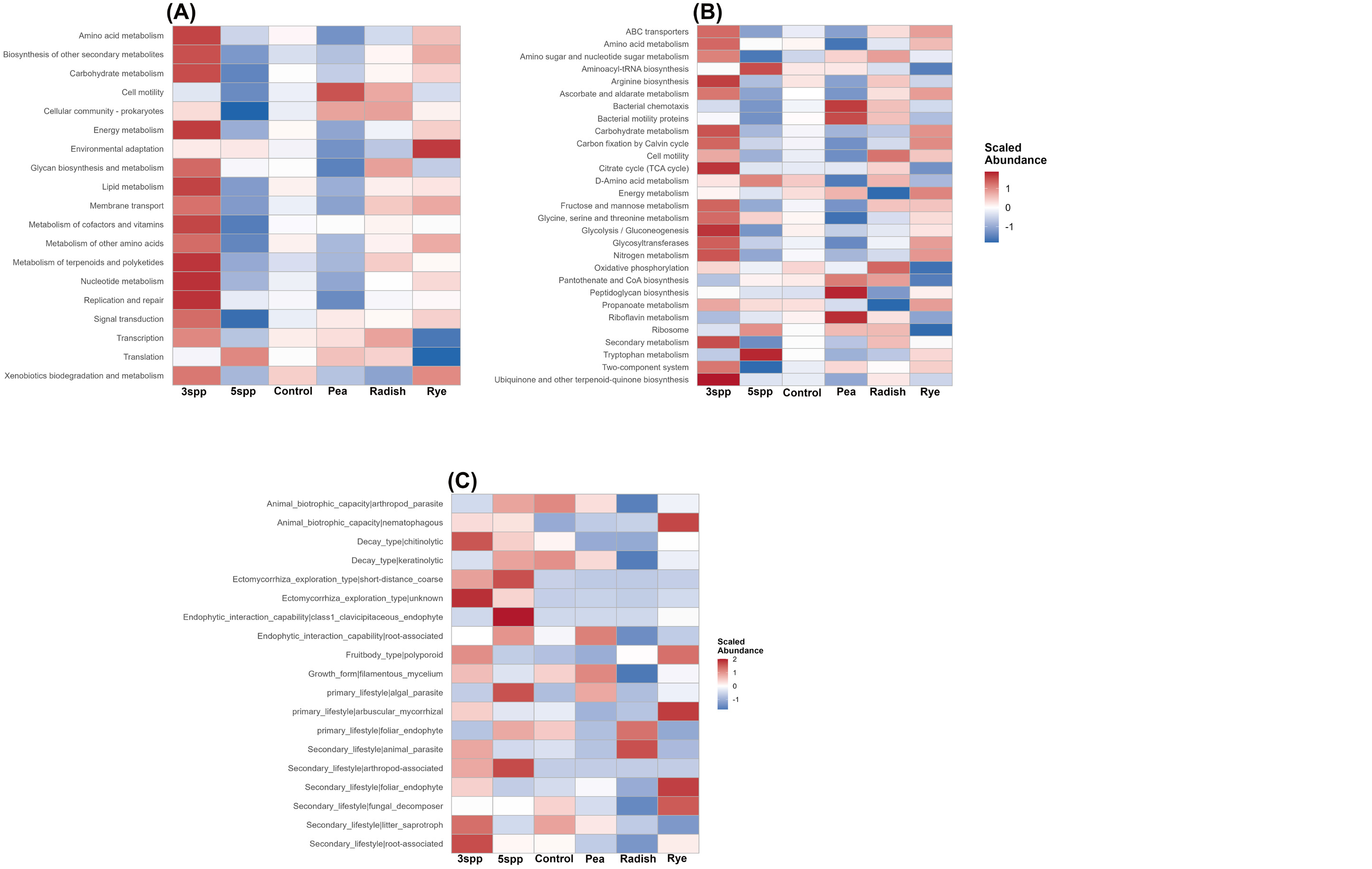
Abundance of predicted bacterial functional categories that were significantly different based on KEGG Category 2 (A) and KEGG Category 3 (B) and fungal functions based on FungalTraits (C). Abundances were first averaged within each CC, then scaled by function (Z-score normalization) across treatments.

At KEGG Level Category 3 (Fig. 4B), finer-scale functional differentiation emerged among treatments. The 3spp. rhizobiome still showed the strongest enrichment across multiple functional categories, including amino acid metabolism, ABC transporters, carbohydrate metabolism, and energy metabolism pathways (citrate cycle, oxidative phosphorylation). The radish rhizobiome exhibited high abundance of ribosome-related functions, secondary metabolism, and oxidative phosphorylation, while showing substantial depletion in carbohydrate metabolism, energy metabolism pathways, nitrogen metabolism, and propanoate metabolism. The rye rhizobiome displayed enrichment in carbohydrate metabolism, carbon fixation by Calvin cycle, energy metabolism, glycosyltransferases, nitrogen metabolism and propionate metabolism, but showed reduction in oxidative phosphorylation, pantothenate and CoA biosynthesis, citrate cycle, riboflavin metabolism and aminoacyl-tRNA biosynthesis. The pea rhizobiome was characterized by elevated peptidoglycan biosynthesis, riboflavin metabolism, bacterial chemotaxis and motility proteins, but showed reductions in amino acid metabolism-related pathways, fructose and mannose metabolism, glycine, serine and threonine metabolism and glycolysis/gluconeogenesis. The 5spp rhizobiome again showed depletion across most functional categories, with particularly strong negative values in secondary metabolism, carbohydrate metabolism, and amino acid metabolism pathways. Most functions for KEGG category 2 and 3 for the control soil were neither enriched or depleted except xenobiotics biodegradation and metabolism which was enriched for KEGG category 2.

Fungal functional trait analysis (Fig. 4C) also revealed distinct predicted ecological strategies across CCs. The 3spp and 5spp rhizobiomes both exhibited elevated abundance of many fungal functions, including ectomycorrhizal exploration types and decay type (chitinolytic). In contrast, all monocultures had more reductions in fungal functions than enrichment of functions. The radish rhizobiome showed increased abundance only in the foliar endophyte and the animal parasite. The pea rhizobiome showed elevation in filamentous mycelium, endophytic interaction capability (root-associated) and algal parasite. Rye monoculture exhibited strong enrichment in animal biotrophic capacity (nematophagous), secondary lifestyle (foliar endophyte and fungal decomposer), arbuscular mycorrhizal, and fruitbody type (polyporoid). Unlike the bacteria functions, the fungal functions in the control soil had more enrichments and depletions such as decay type keratinolytic (enrich), secondary lifestyle litter saprotroph (enrich), and animal biotrophic capacity nematophagous (depleted).

### Differences in Bacterial and Fungal FR

The 3spp rhizobiome had the most significantly different FR KOs, followed by the rye and pea rhizobiomes, based on pairwise comparison analyses, with 3spp, rye and pea having significantly more FR KOs than others (Fig. 5A, Supplemental Table 6). The pairwise comparisons with the most differences were 3spp vs pea and 5spp vs pea. Overall, the control soil had significantly fewer FR KOs compared to all CC rhizobiomes, except for the 5spp rhizobiome. Key KEGG Category 3 functions included ABC transporters, nitrogen metabolism, methane metabolism, quorum sensing, secretion systems, and two-component systems, all of which are linked to microbial activity, nutrient cycling, and environmental response (Supplemental Fig. 4A). Several comparisons highlighted strong directional differences in these processes. The rye rhizobiome compared to the pea rhizobiome had some of the most positive log2 fold changes (more abundant in rye), particularly in carbohydrate metabolism, protein family-related categories, signal transduction, nitrogen metabolism, prokaryotic defense systems, and two-component systems. In contrast, the 5spp rhizobiome had the most negative log2 fold changes (lower than other crops), especially in protein family-related categories, metabolism of cofactors and vitamins, prokaryotic defense systems, and carotenoid biosynthesis. Additionally, the pea vs radish rhizobiome comparison showed mostly negative log2 fold changes for secretion system functions.

**Figure 5:**
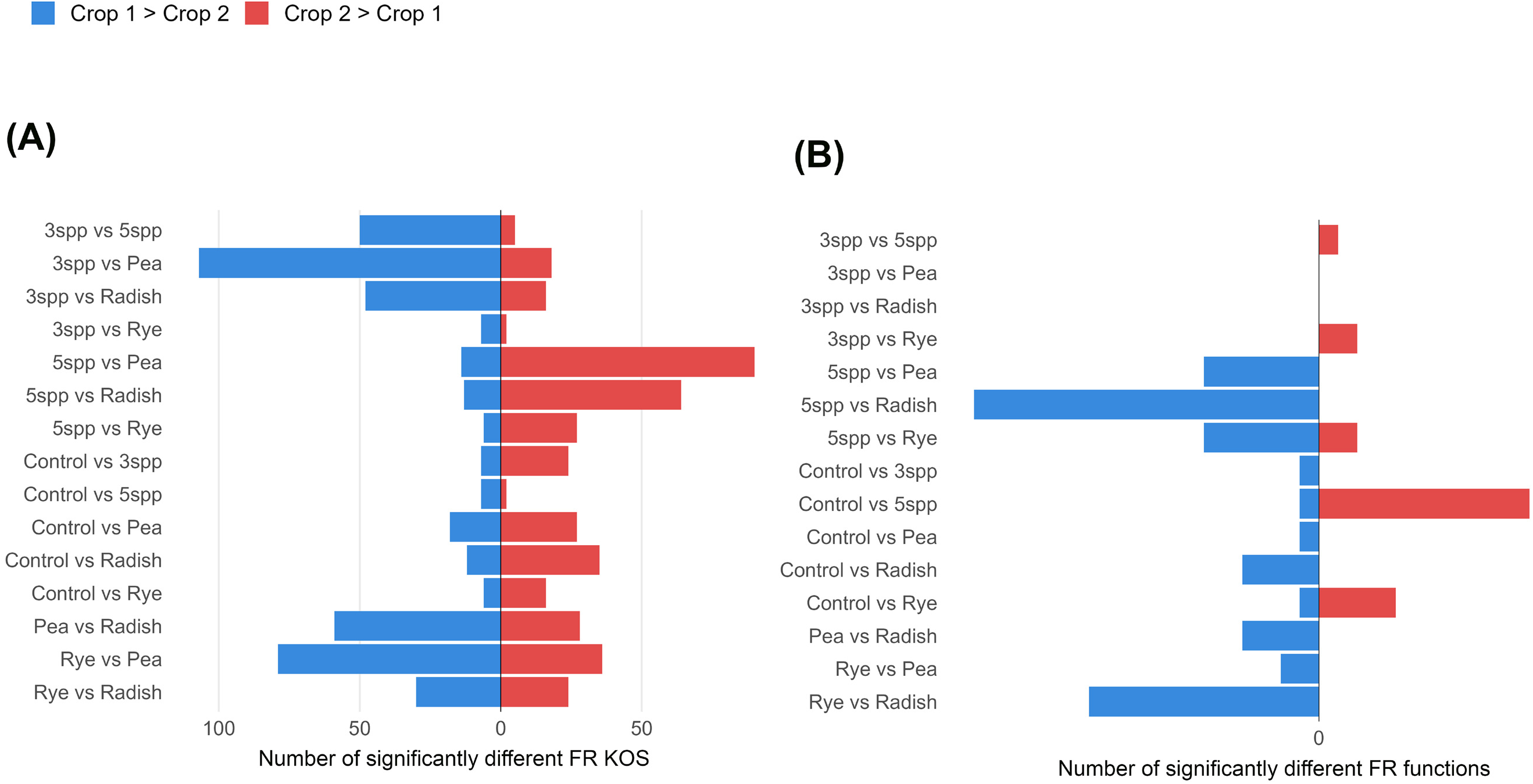
Bacterial FR based on KEGG Orthologs (KOs) (A) and fungal FR based on FungalTraits functions (B). Horizontal bar plots show the number of significantly different Kos or functions (p < 0.05) between pairwise treatment comparisons. Blue bars indicate functions enriched in the first crop listed (Crop 1 > Crop 2), while red bars indicate functions enriched in the second crop (Crop 2 > Crop 1).

For fungal FR, there were relatively few significant differences compared to bacterial FR, likely due to the smaller total predicted functional pool in the FungalTraits database. The highest fungal FR was found in the 5spp rhizobiome, and the lowest in the radish rhizobiome (Fig. 5B; Supplemental Table 6), with the greatest pairwise difference observed between these two treatments. The rye vs radish comparison showed positive log2 fold changes (more in rye) in functions related to decay (white rot), endophytic interactions (root-associated), plant pathogenicity, and ectomycorrhizal lifestyle (Supplemental Fig 4B). The control vs rye comparison showed negative log2 fold changes (more in rye) in functions related to decay substrates, plant pathogenicity, and litter saprotrophy.

## Discussion

Although CCs are known to influence soil microbial communities through root exudation, residue inputs, and alterations in soil physicochemical properties (60, 61), less is understood about how individual CC species and multiple species in mixtures (CC diversity) shape microbial functional potential and FR across both bacteria and fungi. In this study, CC identity and species composition had stronger effects on rhizosphere microbial community composition, predicted functional potential, and functional redundancy than CC species richness alone, as evidenced by the more consistent and pronounced differences among individual CC treatments across taxonomic, functional, and network analyses. While there were limited changes in microbial alpha diversity between treatments, both community composition and predicted functional profiles were strongly structured by specific CC species or CC combinations, indicating that plant functional traits are primary drivers of microbial organization.

Our results partially support Hypothesis 1, as microbial taxonomic composition, functional potential, and FR differed among CC species, particularly between monocultures and mixtures. However, Hypothesis 2 was not supported based on species richness alone, as CC mixtures did not consistently increase microbial diversity or FR relative to monocultures. Instead, results suggest that plant functional composition played a stronger role, with the functionally balanced 3spp mixture showing greater functional potential and FR than the legume-dominated 5spp mixture. Similarly, Hypothesis 3 was not supported when considering diversity alone, as mixtures did not consistently buffer microbial communities from functional or compositional shifts. Contrary to expectations from the diversity–stability hypothesis (62, 63), increasing CC diversity did not consistently enhance microbial function or redundancy, likely due to imbalances in plant functional types and increased competition among microbes. The 3spp rhizobiome, which included a balanced representation of functional groups, showed the highest functional enrichment and redundancy, whereas the 5spp rhizobiome, dominated by legumes, exhibited widespread functional depletion. These results suggest that microbial function is driven more by functional complementarity among plant species than by richness alone. In contrast, monocultures showed strong species-specific effects: rye and pea supported more functionally structured communities, with clearer enrichment of specific metabolic pathways and microbial interactions, while radish supported a simplified rhizobiome characterized by reduced functional diversity, lower network complexity, and more competitive (negative) interactions. In contrast to bacterial patterns, fungal FR was highest in the 5spp rhizobiome, indicating divergent responses of bacterial and fungal communities to CC diversity. Similar patterns were observed in our recent study (64), where species-specific CC effects, particularly in rye rhizobiomes, were more strongly associated with microbial functional stability than plant species richness alone.

### Mixtures differentially shape redundancy-driven stability in soil rhizobiomes

The 3spp rhizobiome showed extensive enrichment across bacterial functional pathways and exhibited the highest bacterial FR among all treatments. Observed enrichments were in key bacterial metabolic processes, including amino acid, carbohydrate, and energy metabolism, as well as transcription, translation, and replication pathways. These patterns suggest that the 3spp rhizobiome hosts metabolically diverse microbial communities with greater growth potential and biosynthetic capacity than those in other treatments. This aligns with the increased diversity and variation of root exudates in multi-species plant systems, offering a broader range of substrates for microbial use (60, 65–67). Besides functional enrichment, the 3spp rhizobiome had the highest degree of bacterial network connectivity, with a larger share of positive than negative interactions. Higher connectivity and positive connections often indicate cooperative or complementary interactions among taxa, which can boost ecosystem functioning and stability (68). The increase in FR further supports the idea that moderate plant diversity fosters microbial communities that are both functionally robust and resilient, with greater overlap in functional potential that may stabilize ecosystem processes despite shifts in community composition (6, 69). However, our previous study involving drought stress (64) demonstrated that these redundancy-associated benefits in a mixed CC rhizobiome may not persist under environmental stress. Under drought conditions, the CC mixture rhizobiome exhibited strong reductions in bacterial FR and increased negative microbial interactions, despite maintaining relatively high taxonomic diversity. This suggests that while balanced mixtures can promote functional complementarity under ambient conditions, drought may constrain the ability of redundant taxa to compensate for functional losses if many taxa respond similarly to stress.

The connection between plant diversity and microbial function was not entirely linear. The 5spp rhizobiome did not always improve microbial functional capacity compared to the 3spp rhizobiome. While there was fairly high functional richness and the most fungal FR, the 5spp rhizobiome had lower bacterial FR, more functional depletion at broader KEGG categories, and a higher percentage of negative bacterial network interactions compared to the 3spp rhizobiome. These patterns suggest that greater plant species richness did not necessarily increase bacterial FR. Instead, the functionally imbalanced 5spp mixture may have promoted niche partitioning or competition among bacteria, leading to more specialized communities with less functional overlap, consistent with ecological theory that diversity effects depend on the balance between complementarity and competition (63, 65, 70). These contrasting patterns between bacterial and fungal communities may reflect fundamental differences in their ecological strategies, where fungi often exhibit higher FR due to broader metabolic capabilities and the ability of multiple taxa to perform similar roles in decomposition and nutrient cycling (69, 71). In addition, fungal co-occurrence networks in this study were less complex and more limited than bacterial networks, suggesting that fungal communities may rely less on interaction structure and more on functional overlap to maintain ecosystem processes.

The lack of a consistent increase in microbial functional capacity in the 5spp rhizobiome may also be explained by differences in plant functional type among CC treatments. While the 3spp rhizobiome included one representative from each major plant functional group (legume, grass, and brassica), the five-species mixture was dominated by three legume species and only one grass and one brassica. This imbalance may have reduced functional complementarity among plants, leading to more uniform resource inputs and fewer distinct ecological niches for microbial communities. In contrast, balanced representation of plant functional types in the three-species system likely promoted a wider diversity of root exudates and soil conditions, supporting greater microbial functional diversity and redundancy. These findings suggest that the functional composition of plant mixtures, rather than species richness alone, plays a critical role in shaping soil microbial community structure and function (63, 65, 70). This interpretation is further supported by our drought experiment (64) where the functionally balanced rye monoculture maintained greater microbial functional stability under drought than the mixed-species treatment. In contrast, the drought-exposed mixture rhizobiome showed increased competitive interactions and widespread downregulation of FR pathways.

Fungal taxonomic differentiation remained relatively limited across treatments, but fungal functional profiles were notably stable, especially in polyculture systems. This separation between taxonomy and function supports the idea of FR in fungal communities, where different taxa can perform similar ecological roles (71). The presence of both mutualistic and saprotrophic groups reinforces the notion that diverse plant systems support multifunctional fungal communities that facilitate nutrient cycling and plant–microbe interactions.

### Monoculture rhizobiomes reflect specialization- and interaction-driven microbial strategies

In contrast to mixture rhizobiomes, monoculture rhizobiomes generally exhibited lower FR and greater functional specialization, particularly in bacterial communities. However, these trends varied across crops, showing that monocultures do not yield a single functional outcome but instead encompass a range of ecological strategies shaped by plant type. Pea rhizobiomes showed an overall decrease in broad functional enrichment, along with an increased presence of motility and root-related traits. These patterns likely reflect strong host-driven selection within the rhizosphere, especially in legume systems where nitrogen inputs and specific root exudates influence microbial community composition (60, 61). Despite overall lower FR compared to CC mixture rhizobiomes, communities supported by peas displayed relatively high bacterial network connectivity and a greater share of positive fungal interactions. This implies a dynamic, rhizosphere-responsive microbial community that may enhance plant–microbe interactions and localized nutrient cycling, even if it lacks the broader functional buffering observed in more diverse systems. However, functions related to nitrogen cycling and symbiosis may be underrepresented in the rhizobiome, as these processes are likely concentrated within root nodules in legume systems rather than distributed across the surrounding soil microbial community (72, 73).

Radish rhizobiomes exhibited a distinct functional profile, marked by enrichment in pathways related to microbial interactions and communication, including secretion systems and quorum sensing. Notably, network analyses indicated that negative interactions largely dominated these communities. This suggests that these pathways may support competitive rather than cooperative processes. Secretion systems and quorum sensing commonly play roles in competition and resource defense (74), indicating that radish fosters a highly interactive but competition-focused microbial environment. While such systems might have reduced FR, they also may enhance microbial turnover or suppress sensitive taxa, reflecting the impact of chemically mediated selection pressures in Brassicaceae rhizospheres (68, 75). Members of the Brassicaceae are generally considered non-arbuscular mycorrhizal hosts and are rich in defense-related glucosinolates, whose hydrolysis products can affect fungal colonization and microbial community assembly; therefore, radish root exudates may impose distinct chemical filters compared with grasses and legumes, contributing to the divergent bacterial and fungal responses observed in the radish rhizobiome (76–78).

On the other hand, rye rhizobiomes often exhibited functional patterns similar to those in the 3spp rhizobiome. The rye rhizobiome showed strong enrichment in pathways linked to nitrogen metabolism, xenobiotic degradation, and secondary metabolism compared to other monoculture rhizobiomes. They also had relatively high FR and fewer differences in KEGG ortholog abundances than in the mixture soil. These outcomes suggest that the rye rhizobiome supports a more specialized yet functionally efficient microbial community capable of maintaining key ecosystem processes. Instead of relying on broad redundancy, rye-associated communities may achieve functional stability through selection of well-suited taxa that can handle complex organic inputs and chemically mediated stress. This aligns with the known influence of rye-derived compounds on soil microbial communities (79, 80). Similar patterns were observed in our drought study (64), where rye rhizobiomes retained stable bacterial FR, network structure, and predicted functional traits under drought conditions. Rye rhizobiomes in the current and previous (64) studies were consistently associated with functionally stable microbial communities, suggesting that grasses may promote drought-resistant microbial assemblages through selective recruitment of stress-tolerant and metabolically efficient taxa.

Network analyses further clarify the differences among monoculture strategies. While monocultures generally showed lower connectivity and a higher proportion of negative interactions compared to the three-species mixture, these patterns varied by crop type. The pea rhizobiome maintained relatively high connectivity, the radish rhizobiome showed competitive (negative) interactions, and the rye rhizobiome displayed a more balanced interaction structure. These distinctions underscore that the stability and function of microbial communities in monocultures rely heavily on plant identity rather than monoculture status alone (68). Beta diversity patterns support these conclusions, especially for fungal communities. While bacterial communities differed overall among treatments without strong pairwise separation, fungal communities exhibited significant pairwise differences, notably between radish and rye. This suggests that fungal communities may be more sensitive to plant identity, although this interpretation should be considered in light of the lower sequencing depth in ITS datasets compared to 16S rRNA gene data, which may influence the resolution of diversity patterns. Bacterial communities, in contrast, may maintain more consistent functionality despite compositional changes, likely due to higher levels of FR (65).

## Conclusions

These results highlight FR as an important mechanism connecting plant diversity to microbial ecosystem function. However, redundancy is not the only way to assess soil processes. Mixture rhizobiomes, particularly the 3spp rhizobiome, supported high levels of FR, broad metabolic capacity, and interconnected microbial networks, all of which signal stable and resilient ecosystems. In contrast, monoculture rhizobiomes typically promote functional specialization, with redundancy concentrated within specific ecological functions rather than distributed across a wide range of processes, resulting in lower overall FR but potentially higher redundancy within targeted pathways. Importantly, the rye rhizobiome likely serves as an intermediate strategy, achieving functional outcomes similar to those in mixture rhizobiomes through functional specialization with redundancy concentrated in specific pathways rather than distributed across a wide range of processes. This suggests that ecosystem functioning can be upheld by various microbial strategies, including stability through FR in diverse systems and efficiency through specialization in more selective environments. These findings align with broader ecological theories, indicating that biodiversity effects arise from both functional complementarity and selection processes (6, 63, 69). In contrast to bacterial patterns, fungal FR was highest in the 5spp rhizobiome, and fungal communities generally showed stronger compositional differentiation but more stable functional profiles, indicating that bacterial and fungal communities respond differently to CC diversity and functional composition.

Overall, this study shows that CC identity and diversity shape soil microbial communities in complex ways that go beyond simple diversity–function relationships. While increasing plant diversity generally enhances microbial FR and interaction networks, some monocultures, especially rye, can achieve similar functional capacity using different ecological strategies. Importantly, comparisons with our drought experiment (64) indicate that these strategies are context dependent. While balanced mixtures promoted high bacterial FR under ambient field conditions in this study, drought conditions reduced redundancy and increased competitive restructuring within mixed rhizobiomes. In contrast, rye monocultures consistently maintained stable functional organization across studies, suggesting that some monocultures may provide greater microbial resilience under environmental stress than more diverse but functionally imbalanced mixtures. These insights emphasize the need to consider both diversity and plant identity when designing CC systems to enhance soil health and ecosystem resilience.

## Acknowledgements

This research used resources on the Palmetto Cluster at Clemson University under National Science Foundation awards MRI 1228312, II NEW 1405767, MRI 1725573, and MRI 2018069. The views expressed in this article do not necessarily represent the views of NSF or the United States government. The authors declare no conflict of interest.

## Supplemental Table Legends

Supplemental Table 1. Summary of Bray–Curtis and UniFrac pairwise distance comparisons used to evaluate whether technical replicates from the same plot were more dissimilar than samples from different plots within the same CC treatment. Pair types include technical replicate pairs from the same plot and same-cover-crop pairs from different plots. Values include the number of pairwise comparisons, mean distance, median distance, and standard deviation.

Supplemental Table 2. Summary of nutrient and soil property comparisons among technical replicate pairs within each CC treatment. For each CC and soil variable, the table reports the number of replicate pairs, Wilcoxon test statistic, raw p-value, Benjamini–Hochberg adjusted p-value, and significance classification. Soil variables include pH, phosphorus, potassium, calcium, magnesium, zinc, manganese, copper, boron, sodium, and nitrate-nitrogen.

Supplemental Table 3. ANCOM-BC2 results identifying differentially abundant bacterial and fungal taxa among cover crop rhizobiomes. Separate worksheets are provided for each dataset. Tables report estimated log₂ fold changes, test statistics, raw and false discovery rate (FDR)-adjusted P values, and significance taxa for pairwise comparisons among cover crop rhizobiomes.

Supplemental Table 4. Co-occurrence network edges for bacterial and fungal ASVs across CC treatments. Edge tables include node identities, correlation direction, edge weight, CC treatment, and taxonomic assignments for each connected node. The bacterial crop-level summary sheet reports the number of positive and negative ASV–ASV connections among phylum pairs within each CC treatment.

Supplemental Table 5. Weighted nearest sequenced taxon index values generated from PICRUSt2 for each bacterial sample. Lower weighted NSTI values indicate closer representation of ASVs by sequenced reference genomes and greater confidence in predicted functional profiles. Maximum and minimum NSTI values across samples are also reported.

Supplemental Table 6. Summary of pairwise comparisons of predicted bacterial and fungal FR among cover crop rhizobiomes. Separate worksheets are provided for bacterial KEGG orthologs (KOs) and fungal FungalTraits functions. For each pairwise comparison, the tables report the total number of predicted KOs or traits evaluated and the number of functions that were significantly enriched in Crop 1 or Crop 2 at *P* < 0.01 and *P* < 0.05.

Supplemental Figure 1: Distance-based Redundancy Analysis (dbRDA) of bacterial communities and fungal communities for TOC, TON, C:N ratio and enzymes (B). Arrows indicate soil chemical variables associated with shifts in community composition. No variables were significantly different.

Supplemental Figure 2. Relative abundance heatmaps of top 10 bacterial (A) and fungal (B) communities across cover crop treatments. Heatmaps display the relative abundance of top 10 families with the associated phylum in parentheses identified in soil samples from different cover crop treatments: 3spp (turquoise), 5spp (red), Radish (green), Pea (pink), Control (purple), and Rye (yellow). Color intensity represents the relative abundance (scale bar, right of each panel), with darker shades of blue indicating higher abundance. Each column represents an individual soil sample, and each row corresponds to a taxonomic group.

Supplemental Figure 3: Barcharts showing the number of positive and negative network connections for bacterial (A) and fungal (B) community comparisons.

Supplemental Figure 4: KEGG level 2 (A) and KEGG level 3 (B) functional categories showing significantly differentially abundant KOs between pairwise crop comparisons. FungalTraits functional categories displaying differential abundance across the same comparisons (C). Each tile represents the log2 fold change in abundance for a given function or trait, with colors indicating direction and magnitude (blue/purple = higher in Crop 1; orange/red = higher in Crop 2). Only functions with statistically significant differences (p < 0.05) and related to soil microbial communities are shown, highlighting shifts in microbial functional composition and trait distributions across cropping systems.

